# Imputation Accuracy Across Global Human Populations

**DOI:** 10.1101/2023.05.22.541241

**Authors:** Jordan L. Cahoon, Xinyue Rui, Echo Tang, Christopher Simons, Jalen Langie, Minhui Chen, Ying-Chu Lo, Charleston W. K. Chiang

## Abstract

Genotype imputation is now fundamental for genome-wide association studies but lacks fairness due to the underrepresentation of populations with non-European ancestries. The state-of-the-art imputation reference panel released by the Trans-Omics for Precision Medicine (TOPMed) initiative contains a substantial number of admixed African-ancestry and Hispanic/Latino samples to impute these populations with nearly the same accuracy as European-ancestry cohorts. However, imputation for populations primarily residing outside of North America may still fall short in performance due to persisting underrepresentation. To illustrate this point, we curated genome-wide array data from 23 publications published between 2008 to 2021. In total, we imputed over 43k individuals across 123 populations around the world. We identified a number of populations where imputation accuracy paled in comparison to that of European-ancestry populations. For instance, the mean imputation r-squared (Rsq) for 1-5% alleles in Saudi Arabians (N=1061), Vietnamese (N=1264), Thai (N=2435), and Papua New Guineans (N=776) were 0.79, 0.78, 0.76, and 0.62, respectively. In contrast, the mean Rsq ranged from 0.90 to 0.93 for comparable European populations matched in sample size and SNP content. Outside of Africa and Latin America, Rsq appeared to decrease as genetic distances to European reference increased, as predicted. Further analysis using sequencing data as ground truth suggested that imputation software may over-estimate imputation accuracy for non-European populations than European populations, suggesting further disparity between populations. Using 1496 whole genome sequenced individuals from Taiwan Biobank as a reference, we also assessed a strategy to improve imputation for non-European populations with meta-imputation, which can combine results from TOPMed with smaller population-specific reference panels. We found that meta-imputation in this design did not improve Rsq genome-wide. Taken together, our analysis suggests that with the current size of alternative reference panels, meta-imputation alone cannot improve imputation efficacy for underrepresented cohorts and we must ultimately strive to increase diversity and size to promote equity within genetics research.

## Introduction

The genome-wide association study (GWAS) is a powerful tool that enables researchers to gain insight into disease mechanisms, discover potential drug targets, and forecast an individual’s risk for health concerns in preventive medicine. Within the past decade, genotype imputation has become an essential component in GWAS. By predicting unobserved genotypes based on a reference panel of externally sequenced individuals, imputation increases marker density, improves the statistical power of GWAS, and enables large-scale meta-analysis across cohorts^1^. Importantly, the size, data quality, and haplotype diversity of imputation reference panels determine the accuracy of genotype imputation^2^. Target populations more genetically related to the reference populations will have better imputation accuracy compared to those more distantly related. Hence, better representation in imputation reference panels will improve the number of diverse populations that can benefit from genotype imputation and GWAS.

In the mid-to late-2000s, one of the first reference panels widely used for genotype imputation in GWAS was based on the International HapMap Project (HapMap), which contained ∼2 Million SNPs discovered through array genotyping and targeted resequencing of three global populations (African-, European-, and East Asian-ancestries)^3^. By the early-to mid-2010s, the sequenced individuals from the 1000 Genomes Project (1KG) formed the basis of an imputation reference panel still used today. However, as GWAS cohorts and consortiums focused on increasing sample sizes among European-ancestry cohorts, so did the large-scale imputation reference panels^4–6^. These Euro-centric reference panels (e.g. ref ^7^) disproportionately performed worse for non-European cohorts. In 2020, the state-of-the-art imputation reference panel released by the Trans-Omics for Precision Medicine (TOPMed)^8^ made an advance towards improving the diversity of reference haplotypes. Though still containing a majority of European-ancestry individuals, the TOPMed panel included a substantial number of admixed African and Hispanic/Latino individuals. As a result, these populations are now much better imputed, with nearly the same efficacy as European-ancestry cohorts. However, previous evaluation of imputation were restricted to reference panels of limited diversity or size, or to target cohorts limited to geographical locale or ancestries^1,3,8–13^. Although imputation is now easier to perform given the publicly accessible imputation server, it remains unclear how well the state-of-the-art imputation reference panel performs for diverse populations across the globe wishing to perform GWAS.

As we push the frontier of genomic medicine to the global scale, it is necessary to be inclusive across diverse populations^9,10^. State-of-the-art imputation reference panels are still largely based on whole genome sequencing data from populations in North America and Europe. We anticipate the TOPMed Reference Panel may fall short in performance due to persisting underrepresentation of diverse global populations, particularly for ethnic minorities residing outside of North America and Europe. To illustrate this point, we curated and evaluated the imputation accuracy across 123 populations worldwide. Because cohort recruitments and GWAS are often conducted in (potentially arbitrarily defined) discrete populations^14^ (though see ref. ^15^) and for the ease of interpreting imputation accuracy metrics, our analysis necessitated defining and assigning descriptive labels to populations across the globe. We recognize that human variations exist on a continuum across space. In this study, we focused on identifying broad patterns across geographical space, rather than implicating specific populations; when we did represent results from a single or a few populations, we did so as illustrative examples.

## Methods

### Dataset Collection

We gathered genotyping array datasets from 23 studies with publication dates ranging from 2008-2022 (**Figure 1**). These represented 14 different genotyping array platforms and 43,507 individuals. These datasets were either publicly available, accessed through online databases such as dbGaP, or requested from the original author of the publication (**Supplemental Table 1**). Before data processing, all genotype and variant information was converted to PLINK Binary file format, and genomic coordinates were converted to GRCh38 using liftOver.

**Figure 1:**
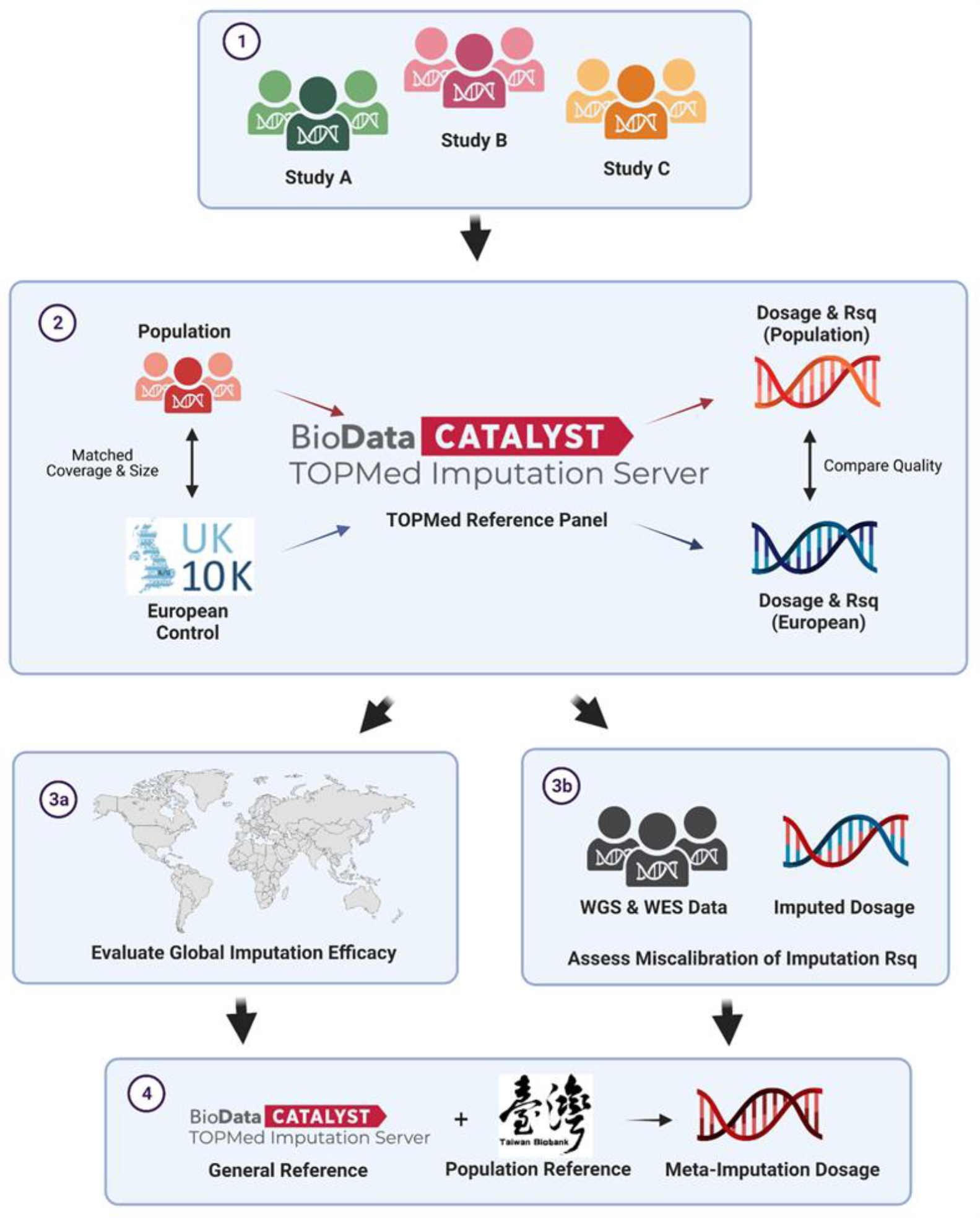
Study Design Overview. (1) We collected and curated genome-wide microarray data from 39 publications published between 2008-2021. In total we imputed 43k individuals and assessed the imputation efficacy in 122 populations across the globe. (2) For each target population we created a matched European control population with the same SNP content and sample size from the UK10K dataset. Both populations were imputed using the TOPMed Imputation Server. (3a) Imputation quality of each target population was evaluated by comparing the mean imputation Rsq in strata of allele frequencies to that of the matched European controls. (3b) For selected populations with available sequencing data, we assessed imputation quality using Pearson’s r2 comparing the imputed dosage with the genotype from sequenced data. (4) We assessed meta-imputation to improve imputation quality by combining imputed dosages from TOPMed and Taiwan Biobank (TWB).

### Quality Control & Population Consolidation

For each dataset, individuals with greater than 5% missingness and variants greater than 2% missingness were removed from the analysis. When genetic ancestry profiles are available from the original publication, we manually grouped populations with similar ancestry profiles into larger populations and then removed populations with less than 20 individuals from downstream analyses. Across all of our studies, we assembled 123 populations with 20 or more individuals for analysis (**Supplemental Tables 1-2**). Once the population to be imputed is defined, we then filtered variants that failed the Hardy-Weinberg equilibrium (p < 10^6^) per population. The number of variants remaining at each step of pre-imputation filtering is provided in **Supplemental Table 3**. Next, the microarray data were converted to variant call format (VCF) files by chromosome for submission to the imputation server. Only autosomal markers were imputed and analyzed. If > 10% of strand flips were identified during the imputation server quality control that prevented imputation from proceeding, then we correct the strand flips by comparing the two alleles for each variant against those recorded in the 1000 Genomes sequencing data. For each population, we estimated latitude and longitude using the Geocoding API feature in the Google Maps Platform and location descriptors in the original sources; latitude and longitude used to generate **Figure 2A** and **Supplemental Figures 1-2, 6** are provided in **Supplemental Table 1**.

**Figure 2:**
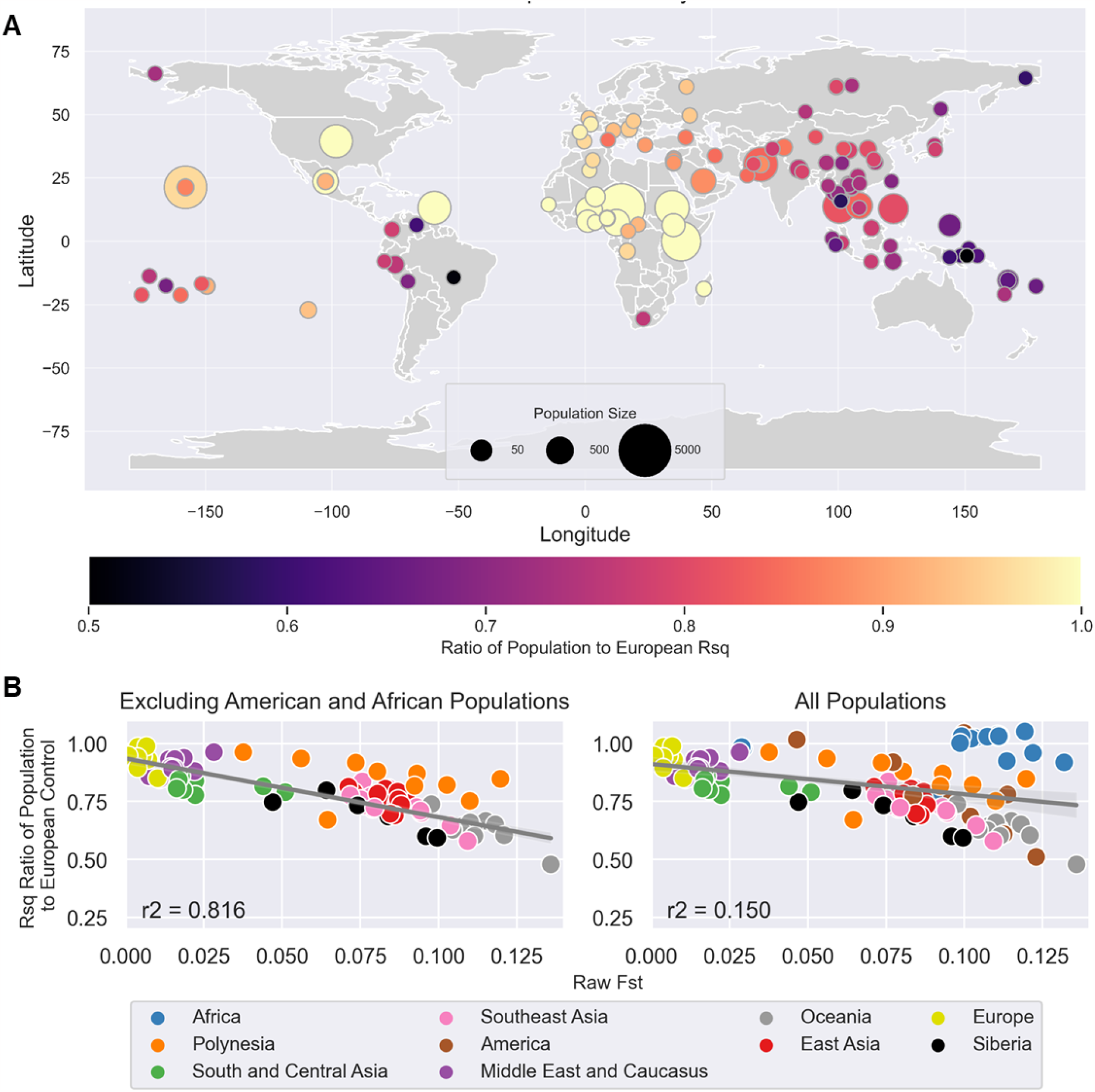
Disparity of TOPMed imputation accuracy across the globe. (A) Ratio of the mean imputation Rsq of 1-5% alleles for every target population to its corresponding matched european control are shown on the world map. Values closer to 1.0 indicate no relative imputation accuracy loss compared to a European cohort matched by sample size and SNP content. Size of the circle scales with the sample size of the dataset that was imputed, and color of the circle scales with imputation accuracy. (B) Relationship between the ratio of mean Rsq and Fst between each target population and its European control. Each population is colored according to its assigned broad geographical region. The left plot excluded African and American populations. The right plot contains all populations analyzed, where notably African-ancestry populations show strong genetic differentiation from Europeans by Fst, but are imputed with high efficacy using TOPMed.

### Matched European-Ancestry Population for Comparisons

For each target population we curated and processed for imputation a European-ancestry population using the UK10K dataset^16^ (European Genome-phenome Archive accession numbers EGAD00001000740 and EGAD00001000741), which contained whole genome sequencing data from the ALSPAC and TwinsUK cohorts. For each target population, we matched the sample size and SNP content in the UK10K dataset in order to account for differences in performance due to array types and for better interpretation of imputation accuracy across different allele frequencies. We extracted randomly matching number of individuals from the UK10K dataset,and extracted all SNPs in the target population passing quality control for imputation and also found in the UK10K dataset. Note that not all SNPs may be found in the UK10K dataset, thus our comparison population for imputation accuracy based on the European-ancestry population necessarily has equal or fewer SNPs used for input of imputation. However, this is expected to lower the quality of imputation for the European-ancestry population and reduce any disparity in imputation accuracy observed, if any.

### TOPMed Imputation and measure of imputation accuracy

Unphased populations were submitted to the TOPMed imputation server for further server-based quality control and imputation with the mixed population setting. The Rsq filtering was turned off and Eagle v2.4 (ref ^17^) phasing was selected. The defaults for the remaining settings were selected.

In general, there are two types of statistics to measure imputation accuracy: statistics produced without reference to true genotypes (*e*.*g*. INFO score, minimac’s R-squared [Rsq]) and statistics that compare imputed to genotyped data (*e*.*g*. LooRsq, concordance rate, Pearson’s squared correlation, or Imputation Quality Score [IQS])^18–21^. In general, we do not have access to true genotypes and thus elected to use minimac’s Rsq as our primary measure of imputation accuracy across the full frequency spectrum, because it is widely used for filtering markers post-imputation for association testing. For each population, including the European controls, the imputation Rsq metric was extracted for each imputed variant from info files produced by the imputation software, Minimac 4. Distributions of Rsq were examined in strata of minor allele frequencies based on the estimated frequency from imputation.

For a subset of populations where publicly available whole genome seqeuncing data were available, we also compute Pearson’s squared correlation, r^2^, as an additional measure of imputation accuracy, and for comparing to the imputation Rsq. We chose to use Pearson’s r2 since it gives a directly interpretable value to relative statistical power in GWAS, and because compared to arguably more advanced metric such as IQS, it is equally well-calibrated for common variants^21^. We assessed populations from the Human Genome Diversity Project^22^ (HGDP) and the African American Sequencing Project from 23andMe (AASP, dbGAP accession number phs001798.v2.p2). For HGDP, we processed the previously released genome-wide array data^23^ as described above for imputation. For the AASP cohort, we extracted the SNPs found on Omni 2.5M array to mimic an array dataset. For overlapping individuals and SNPs between the imputed data and sequencing data, we computed Pearson’s correlation r^2^ between the imputed dosages and the genotypes from sequencing data.

### Fixation Index (Fst) Calculation

Each target population was merged with the matched European-ancestry control from UK10K, and the intersection of variants was kept. Variants were pruned to achieve approximate linkage equilibrium using PLINK^24^ (v1.9, using the --indep-pairwise command with a window size of 50kb, step size 5, and r-squared threshold 0.1). Fst was calculated using the --fst command. Raw Fst score was used to estimate population differentiation.

### Meta-Imputation

To assess meta-imputation, we imputed selected populations with Taiwan BioBank ^25,26^ (TWB) as an imputation reference in-house. Whole genome sequenced individuals from TWB were phased without a reference using Eagle (version 2.4v). Phased genotypes were then converted into the reference file format (.m3vcf files) with Minimac 3 (ref. ^19^). To perform meta-imputation, selected target populations were phased independently without a reference using Eagle 2.4v first (rather than with the TOPMed imputation server), and imputed with TWB imputation reference panel in-house using Minimac4 (ref. ^18^), or on the TOPMed Imputation Server. The imputation results for TWB and TOPmed were meta-imputed with MetaMinimac2^27^. To evaluate the improvement of imputation Rsq at East Asian-enriched alleles, we stratified each meta-imputed variant according to their estimated frequencies from the Genome Aggregation Database (gnomAD) non-Finnish-Europeans and East Asians (release 3.1.1; https://gnomad.broadinstitute.org/)^28^.

## Results

### Persisting disparities in imputation accuracy with the TOPMed reference panel

To identify the current insufficiencies of the TOPMed panel for imputing diverse global populations, we curated genome-wide array data from 23 studies published between 2008-2021 (**Methods, Figure 1**). In total, we imputed over 43k individuals across 123 populations, spanning multiple array platforms, three orders of magnitudes in sample sizes, and ten major geographical regions around the globe (**Supplemental Figures 1-2, Supplemental Tables 1-2**). As the true genotypes across the entire allele frequency spectrum are not available for vast majority of these populations, we assessed the imputation accuracy of each population by comparing the distribution of imputation Rsq, stratified by bins of minor allele frequencies, to that of a European cohort (the UK10K sequencing data)^29^ subsampled to match in sample size and array SNP content (**Methods; Figure 1**). By matching to the UK10K data in sample size and SNP content, we will be able to account for differences in imputation accuracy due to different array types and SNP density, and be able to interpret the relative imputation accuracy between two populations within the same frequency bin. Indeed, we generally observed that arrays with higher marker density tend to perform better in imputation compared to those with lower marker density, and that 5-50% alleles (**Supplemental Figure 3B**). are imputed more accurately than 1-5% alleles variants (**Supplemental Figure 3A**).

**Figure 3:**
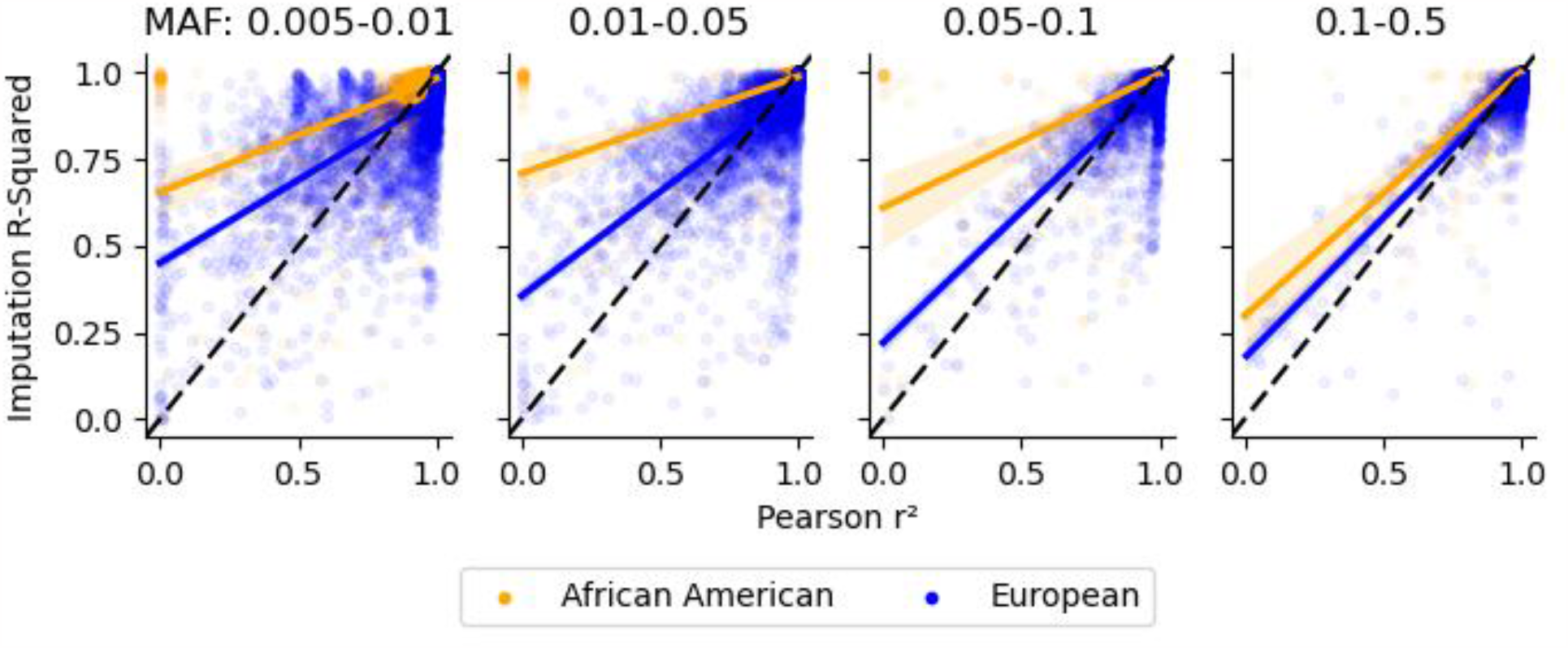
Imputation Rsq is an overly optimistic metric of imputation efficacy. The relationship between Pearson’s r2 and Minimac4-Produced Imputation Rsq. A random set of 5,000 variants in each strata of minor allele frequency were sampled from 23andMe’s African American Sequencing Project data (N=2269) and Europeans from the Human Genome Diversity Project (N=155). The dash line denotes unity, while the solid lines are fitted linear models. Comparing the Imputation Rsq to Pearson’s r^2^ for the same set of variants, Imputation Rsq appears to overestimate imputation quality on average. Imputations of African American samples experienced a higher inflation of Imputation Rsq compared to that of Europeans across all minor allele frequency bins.

Most importantly, the imputation of the non-European populations tends to be worse than that of the matched European populations, particularly for alleles with less than 5% frequency (**Supplemental Figure 4, Supplemental Table 4**). To identify broad patterns across the globe, we visualized the ratio of mean Rsq between the target population and the matched European control population on a world map. From these comparisons, populations from Asia (East, Southeast, South), the Middle East, Siberia, Oceania, and the Pacific have the overall lowest imputation accuracy compared to matched controls from Europe, presumably due to underrepresentation of haplotypes from these regions within the TOPMed panel (**Figure 2A**). For instance, the mean Rsq for 1-5% alleles in Saudi Arabians (N=1061)^30^, Vietnamese (N=1264)^31^, Thai (N=2435)^31^, and Papua New Guineans (N=776)^31^ were 0.79, 0.78, 0.76, and 0.62, respectively. In contrast, the mean Rsq ranged from 0.90 to 0.93 for matched European populations (**Figure 2A**). On the other hand, African-ancestry populations as well as some American and Pacific populations imputed relatively well compared to European populations, likely due to their representation, directly or through shared ancestry from recent admixture, in TOPMed (**Figure 2A**). Expectedly, the imputation accuracy for a target population is inversely correlated with the genetic differentiation, as measured by the fixation index (F_ST_), to the matched European populations (**Figure 2B**). The exceptions to this relationship are those populations from Africa or the Americas, which reflects the improved representation of African-ancestry individuals and Latinos in the TOPMed panel (**Figure 2B**). We observed qualitatively similar but less extreme patterns for the imputation accuracy of alleles 5% or greater in frequency (**Supplemental Figure 5**).

**Figure 4:**
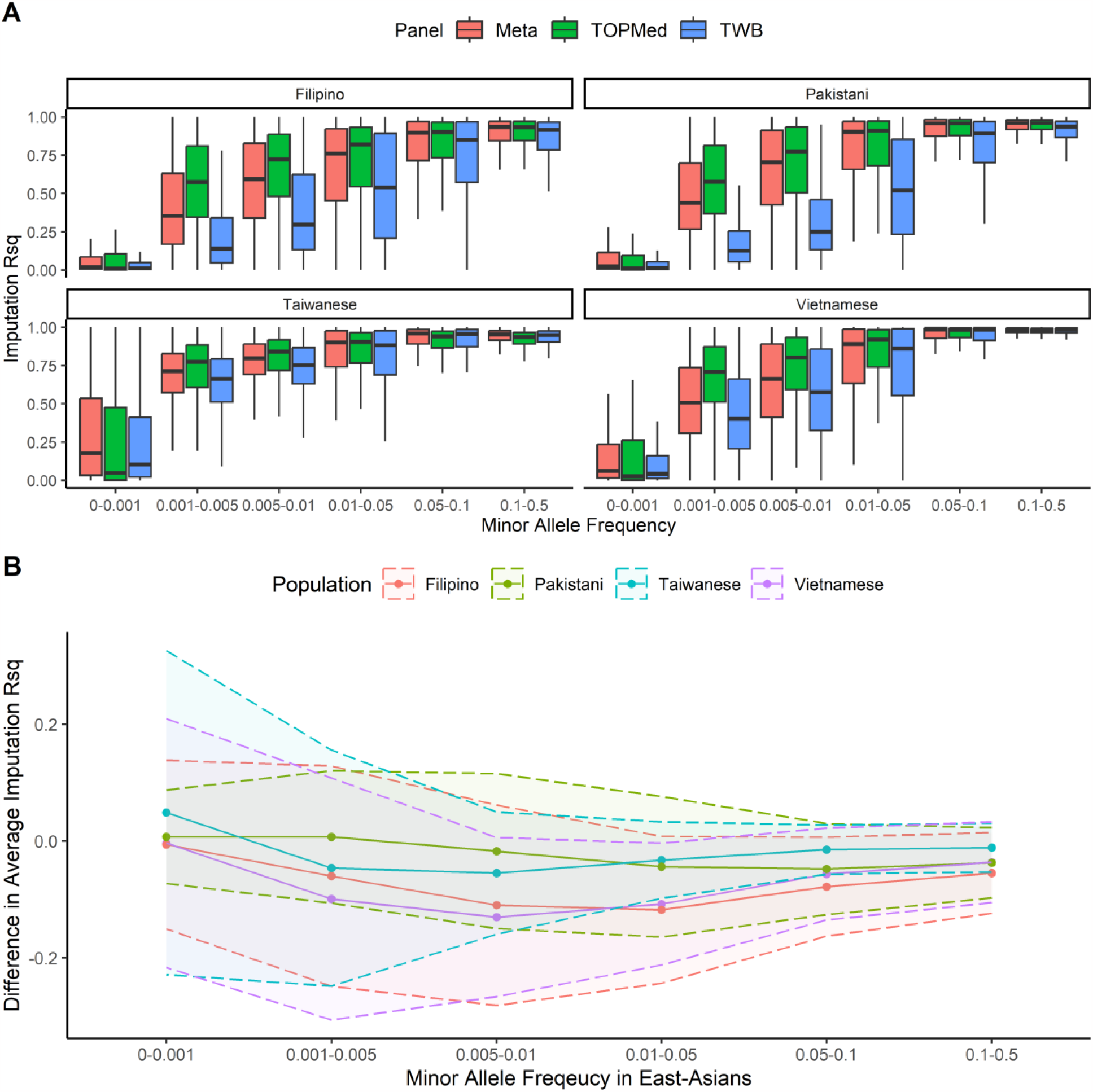
Meta-Imputation does not improve imputation accuracy. (A) Four populations, Filipino (N=1799), Pakistani (N=2493), Taiwanese (N=1999), and Vietnamese (N=1264) were independently phased and imputed with both Taiwan BioBank (TWB) and TOPMed. Imputation results were meta-imputed and the distribution of Imputation Rsq across different strata of allele frequencies are shown for imputation with TOPMed (blue), TWB (red), and meta-imputation (green). Meta-imputation only marginally improves the imputation accuracy as measured by Rsq for common variants, if at all, even in cases when the target (e.g. Taiwanese) is from the same population as one of the references (TWB). (B) Restricted to variants with minor allele frequency less than 0.1% in gnomAD non-Finnish Europeans, meta-imputation combining TOPMed with TWB does not improve the imputation accuracy for variants that are enriched in East Asians, relative to just imputing with TOPMed alone. A genome-wide random sample of 10% of the SNP included in both TWB and TOPMed was taken for analysis. Lines reflect the mean differences between meta-imputation and TOPMed-imputation alone. Shaded bands reflect the standard deviation across variants within the frequency bin.

We developed an interactive website that stores and displays summaries of all of our imputation results for users to access: https://jlcah5.github.io/iaw/map.html. Taken together, the discrepancies in imputation accuracy across the frequency spectrum suggest that some populations across the globe, notably those from Asia, Oceania, and the Pacific, will not reap the full benefits of imputation of a large reference like the TOPMed panel, and suffer a relative loss in statistical power in GWAS even if large-scale recruitment of study cohorts is feasible.

### Inferred gaps in imputation accuracy across populations may be overly optimistic

For many of the populations assessed, we relied on Rsq estimated by the imputation software as a metric of imputation accuracy. However, imputation Rsq is estimated from imputed frequencies without information from observable frequencies, and as a result may misrepresent the true imputation accuracies.^32^. A better estimate of imputation accuracy, particularly in relation to the potential loss of statistical power in GWAS, is based on Pearson’s correlation coefficient (r^2^), computed between the imputed dosage and the observed genotype. For selected populations where we have access to publicly available high-coverage genome sequencing data (**Supplemental Table 5**), we used Pearson’s r^2^ to measure imputation accuracy.

In general, we found a qualitatively similar pattern using Pearson’s r^2^: common variants are imputed better than rare variants, and European-ancestry populations are imputed better than other non-European populations (**Supplemental Figure 6**). However, when comparing the two measures of imputation accuracy (imputation Rsq vs. Pearson’s r^2^), we observed that imputation Rsq is inflated compared to Pearson’s r^2^. While the inflation is evident across the frequency spectrum, it is more inflated among the rarer alleles (**Figure 3, Supplemental Table 6**). Importantly, imputation Rsq appears to be biased upwards in non-European populations than European populations compared to Pearson’s r^2^ (**Figure 3**). For instance, mean imputation Rsq for 10-50% frequency alleles genome-wide were similar between African-ancestry populations and their matched European counterpart (**Figure 2A**), suggesting similar imputation accuracies. But for an allele between 10-50% frequency in Europeans with an estimated imputation Rsq of 0.5, the expected Pearson’s r^2^ is a relatively well-calibrated 0.42. In contrast, for an allele in the same frequency range and estimated Rsq in African Americans, it would have an expected Pearson’s r^2^ of 0.23, a 45% relative decrease in power when testing the association of this imputed variant with phenotype. We also observed excess inflation across the frequency spectrum for other populations from the Human Genome Diversity Project that we assessed (**Supplemental Figures 7-15**). Therefore, while we have shown reduced efficacy when imputing diverse populations using the current TOPMed imputation resources, our global assessment using Rsq may underestimate the degree to which disparities in imputation accuracy exist. Our results here suggest that true imputation accuracy, as well as the potential loss in statistical power in GWAS, may be even lower for non-European populations.

### Meta-Imputation does not improve imputation efficacy for underrepresented cohorts

Populations in key regions around the world are imputed with comparably lower quality using the TOPMed reference panel due to underrepresentation within the imputation reference. We speculate that one approach to improve imputation accuracy and circumvent the issue of underrepresentation in the TOPMed panel is meta-imputation. Meta-imputation is a tool that combines the imputed dosages after imputation across multiple panels using weights based on empirically estimated imputation performance at genotyped sites^27^. Specifically, one could combine the imputation result from the TOPMed reference panel with a smaller, regional or population-specific imputation panel to improve the imputation accuracy, while still leveraging the large size of the TOPMed panel for better imputation of common variants.

We tested the efficacy of meta-imputation across the genome for the Filipinos (N = 1,799)^33^, Pakistani (N=2493)^34^, Taiwanese (N=10,000)^25^, and Vietnamese (N = 1,264)^31^, chosen for their relatively large sample size and poor imputation using TOPMed alone (**Figure 2A**). We meta-imputed the TOPMed-imputed results with that of a reference panel constructed from whole genome sequenced individuals from the Taiwan BioBank (TWB; N = 1,496)^25,26^. Because many Taiwanese individuals are of Southern Chinese origin, and aboriginal Taiwanese may be related to their ancestors that expanded across the Pacifics^35,36^, there may be some benefit to using TWB for the meta-imputation of Southeast Asian individuals. In total, 20,698,795 variants were found in both the TOPMed and TWB panels. In general, imputation accuracy with the TWB panel was substantially poorer than that with the TOPMed panel, even for Taiwanese individuals (**Figure 4A**). This is likely due to the sheer sample size difference of the TOPMed and TWB imputation reference panels. As a result, meta-imputation between TOPMed and TWB panels overall rarely improved imputation accuracy, as quantified by imputation Rsq, and only marginally improved Rsq for common alleles, if at all (**Figure 4A**). For instance, for 1-5% alleles, meta-imputation achieved a mean Rsq of 0.75, 0.64, 0.75, and 0.74, for Vietnamese, Filipino, Pakistani, and Taiwanese, respectively. This is comparable to the mean Rsq from imputing with the TOPMed panel alone (0.79, 0.68, 0.76, and 0.73 for the same populations, respectively).

We also examined if alleles enriched in East Asians experienced a boost in imputation accuracy through meta-imputation with a East Asian-specific reference panel, but generally observed no improvement over TOPMed (**Supplemental Figures 16-19**). For example, restricting to a 10% subsample of SNPs genome-wide (∼2M alleles) that were found in less than 0.1% frequency in Genome Aggregation Database (gnomAD)^28^ non-Finnish Europeans but are found in 1-5% in gnomAD East Asians, we observed that Rsq for Vietnamese, Filipino, Pakistani and Taiwanese populations actually decreased by 10.8%, 11.8%, 4.4% and 3.2%, respectively when meta-imputed (**Figure 4B**). Meta-imputation did appear to improve the imputation accuracy for more common alleles compared to TOPMed imputation for the Taiwanese, perhaps reflecting the specificity of the TWB reference panel (**Supplemental Figure 18**), but this did not translate to other South and Southeast Asian populations. We note that while meta-imputation did not generally improve imputation accuracy of population-enriched variants, meta-imputation increased the variant coverage available for analysis. When meta-imputing with the TOPMed (292,145,137 SNPs) and TWB (39,935,638 SNPs) reference panels, there was an addition of 930,696 variants from TWB that would become available for analysis mostly due to extremely rare variants in TWB not found in TOPMed (**Supplemental Figure 20**). Meta-imputing TOPMed and TWB together led to a 6.5% increase in SNP coverage compared to imputing with the TOPMed panel alone.

## Discussion

Genotype imputation is a critical tool that allows us to conduct effective, population-scale association studies. Our work reveals key discrepancies driven by the lack of global representation in the state-of-the-art imputation panel generated by the TOPMed consortium. Across 123 populations and >43,000 individuals, we concluded that imputation accuracy is generally lower for diverse populations across the globe, especially those residing outside of the Americas and Europe, compared to matched European controls. For instance, for 1-5% alleles, populations from East Asia, the Middle East, Oceania, Polynesia, Siberia, Central and South Asia, and Southeast Asia are imputed with average reductions of 6.5-42% in imputation Rsq compared to controls (**Figure 2A, Supplemental Table 4**). The differences in imputation accuracy are driven by the under-representation of non-European and non-American populations in TOPMed.

We note that while our analysis covered many global regions, there were a number of areas that we could not adequately gather data comprehensively. For example, our Sub-Saharan African populations are dominated by Western and Eastern Africans. While our results showed that African populations are imputed with similar quality to the matched European populations (based on Rsq), this is likely reflecting the predominant West African ancestries that are found in the African American populations in the TOPMed reference panel^37^. The same level of imputation accuracy may not generalize to the Southern African regions^11^. In fact, imputation Rsq for South Africans is 0.28 lower than that of their European controls for 5-50% alleles **(Supplemental Table 1, Supplemental Figures 4-5)**. More African datasets outside of West Africa are emerging^11–13^ and future assessments are needed to better characterize the generalizability of current reference panels to the rest of the African continent.

It is clear from our study that we must strive for sequencing larger cohorts of individuals from diverse populations, particularly if meta-imputation facilitates future imputations at large scale. Meta-imputation of the 1000 Genomes Project and the Haplotype Reference Consortium (HRC) imputation panels were previously shown to have nearly the same imputation accuracy as combining both imputation panels and better imputation accuracy than imputing with either panel alone^27^. In our assessment, one major challenge with the current meta-imputation framework is the drastic size differences between TOPMed and population-specific panels such as TWB, on top of potential differences in the generation of the sequencing data. The TOPMed reference panel contains approximately 65-fold more individuals and nearly 7-fold the number of sites as TWB. Only ∼7% and ∼52% of variants from TOPMed and TWB, respectively, were found commonly on both panels and available for meta-imputation. As a result, meta-imputation may not consistently improve imputation accuracy when the TOPMed reference panel is involved, particularly for rare variants as these are less likely to be shared among imputation panels. As more whole genome sequencing data become available, particularly from diverse regions around the world, we expect the benefit of meta-imputation to become more apparent for non-European populations.

Looking ahead, advances in three specific areas related to imputation and the construction of future reference panels are needed to help drive inclusivity efforts in human genetic studies. First, current large scale resequencing projects tend to be performed on cohorts recruited as part of large epidemiological studies. These study participants tend to be sampled from major metropolitan areas, sometimes even residing outside of the country of origin. Hence, these larger cohorts may not fully represent the haplotypic diversity of all local populations in their respective regions. Oftentimes these local populations are in greater needs to benefit from the fruit of advances in genomic medicine. Importantly, individuals with higher components of indigenous ancestries tend to be imputed with lower quality. For example, Native Hawaiian individuals with elevated levels of Polynesian ancestries^38^ were imputed with ∼0.1 lower Rsq in 1-5% alleles compared to randomly sampled admixed Native Hawaiians^35^ (**Figure 2A, Supplementary Table 1, Supplemental Figure 4**). Therefore, to increase diversity and representation of imputation references, we must specifically and ethically sample ethnic minorities from diverse regions around the globe to complement the epidemiological cohorts from these regions.

Second, meta-imputation currently requires specific file format and information (empirical dosage) that is only output by the imputation software Minimac^18^. As some population reference panels are hosted on different imputation servers with alternative algorithms, we are not yet able to meta-impute against all possible reference panels. For instance, it has been shown that TOPMed reference panel, along with the African Genome Resource^39^, appear to impute Sub-Saharan populations with the highest efficacy^11^. But because the African Genome Resource is hosted on Sanger Imputation Service with a different imputation software using the Positional Burrows Wheeler Transform algorithm^40^, it cannot be meta-imputed with TOPMed. A consensus within the field to output standardized imputation information is thus required if the individual level data for imputation references cannot be shared publicly.

Third, sequencing data of large reference panels need to be more accessible and data processing should be more standardized. Sequencing studies of diverse populations have been emerging, resulting in a number of population-specific imputation reference panels. For instance, a reference panel for African Americans enriched with Sub-Saharan ancestry^41^ (N=2294) showed similar performance as TOPMed and significantly outperformed other imputation panels, such as 1KG and HRC, for imputing African American individuals^42^. While regional panels at their current sample sizes may not improve overall imputation compared to TOPMed, a number of panels, including the Japanese population reference panel (N=1070)^43^, the China Metabolic Analytics Project (N=10155)^44^, the Mexico City Prospective Study (N=9950)^45^, a reference panel containing primarily Qatari individuals, a reference panel containing East Asian individuals (N= 14393)^46^ and Arab individuals (N=12432)^47^, and a reference panel of Ashkenazi Jewish individuals (N=128)^48^, have demonstrated that high coverage sequencing and the construction of population-specific panels have the potential to be invaluable for identifying rare, disease-causing variants. Many of these risk alleles may be novel, population-specific, and not included in other resources. However, currently access to these data are generally strictly controlled, and even if hosted in an imputation server it may be hosted with variable QC pipelines and output information, potentially compromising the effectiveness of meta-imputation or preventing a uniform assessment of imputation accuracy or meta-imputation framework.

It is important that we maximize the number of people who can benefit from large-scale association studies – understanding the representativeness and performance of the current state-of-the-art imputation reference panel is an important first step. As global sequencing efforts progress, we expect the release of more regional imputation panels will initiate a wave of studies identifying population-specific genetic association to complex traits and diseases. It is imperative to host these future panels on publicly accessible imputation servers, ideally with harmonized imputation output, if the individual level data cannot be shared. We must continue to sequence diverse individuals and strive toward more representation in our scientific resources in order to achieve equity in genetic research.

## Acknowledgements

The authors would like to thank Xin Sheng and Amelia Marvit for comments and feedback of this manuscript. This work was supported by funding from the National Institutes of Health (R35GM142783 to C.W.K.C.). J.L.C. and X.R. were supported by the Viterbi Merit Fellowship and the Provost Fellowship, respectively, through the University of Southern California. In addition, this study benefited from a number of scientists who kindly shared their data for analysis, including Georgi Hudjasov, Phillip Endicott, Meshari Alwazae, Fowzan Alkuraya, Dang Liu, Mark Stoneking, and David Reich; the original publications are cited within the text. This study also accessed several datasets through online repositories such as dbGaP. For these studies, we acknowledge the use of:

1. Data generated by MalariaGEN. A full list of the investigators who contributed to the generation of the data is available from www.MalariaGEN.net. Funding for this project was provided by Wellcome Trust (WT077383/Z/05/Z) and the Bill & Melinda Gates Foundation through the Foundation of the National Institutes of Health (566) as part of the Grand Challenges in Global Health Initiative.
2. Data generated by the Pacific Islands Rheumatic Heart Disease Genetics Network. A full list of the investigators who contributed to the generation of the data is available from www.rhdgenetics.net. Funding for the project was provided by the British Heart Foundation (PG/14/26/30509; www.bhf.org.uk), the Medical Research Council UK (Fellowship G1100449; www.mrc.ac.uk), and the British Medical Association (Josephine Lansdell Grant; www.bma.org.uk) and cite the relevant primary Network publication (details of which can be found at www.rhdgenetics.net).
3. Data generated by the Genetic Risk to Stroke in Smokers and Non-Smokers in Multiple Ethnic Groups study, funding for which was provided through the NIH Genes, Environment and Health Initiative [GEI] (U01HG004436). The Genetic Risk to Stroke in Smokers and Non-Smokers in Multiple Ethnic Groups study is one of the genome-wide association studies funded as part of the Gene Environment Association Studies (GENEVA) under GEI. Assistance with phenotype harmonization and genotype cleaning, as well as with general study coordination, was provided by the GENEVA Coordinating Center (U01 HG004446). Assistance with data cleaning was provided by the National Center for Biotechnology Information. Funding support for genotyping, which was performed at the Johns Hopkins University Center for Inherited Disease Research, was provided by the NIH GEI (U01HG004438) and the NIH contract “High throughput genotyping for studying the genetic contributions to human disease”(HHSN268200782096C). Support for the fieldwork in Pakistan was funded from the NINDS (R21NS064908) and support for genotyping of common controls was from the Wellcome Trust.
4. Data generated by the Diabetes in Mexico Study SIGMA Exome Sequencing Project, which was supported by the Carlos Slim Foundation. The data from the Diabetes in Mexico Study SIGMA Exome Sequencing Project reported here were supplied by the Broad Institute and National Institute of Genomic Medicine (INMEGEN). This manuscript was not prepared in collaboration with Investigators of the Diabetes in Mexico Study SIGMA Exome Sequencing Project study and does not necessarily reflect the opinions or views of the Diabetes in Mexico Study SIGMA Exome Sequencing Project study, or the Carlos Slim Foundation.
5. Data generated by the Stanford University Global Reference Panel data. Funding support for the Population Architecture Using Genomics and Epidemiology (PAGE) Global Reference Panel was provided through the National Human Genome Research Institute (U01HG007417). Assistance with phenotype harmonization, SNP selection, data cleaning, meta-analyses, data management and dissemination, and general study coordination, was provided by the PAGE Coordinating Center (U01HG007419, U01HG004801 and its NHGRI ARRA supplement). The datasets used for the analyses described in this manuscript were obtained from dbGaP.
6. Data generated by the African American Sequencing Project, which was supported by the National Human Genome Research Institute of the National Institutes of Health (grants R44HG009460 and R43HG009090). We thank the research participants and employees of 23andMe for contributing to this study.
7. Data generated by the UK10K Consortium, derived from samples from the TWINSUK and ALSPAC cohorts (EGAD00001000740 and EGAD00001000741). A full list of the investigators who contributed to the generation of the data is available from www.UK10K.org. Funding for UK10K was provided by the Wellcome Trust under award WT091310.

Finally, computation for this work was supported by the University of Southern California’s Center for Advanced Research Computing (CARC; https://carc.usc.edu).

## Notes

https://jlcah5.github.io/iaw/map.html

### Competing Interest Statement

The authors have declared no competing interest.

### Summary of Updates

Section on Meta-Imputation was updated with new results.

