## Supplemental Figures for "Imputation Accuracy Across Global Human Populations"

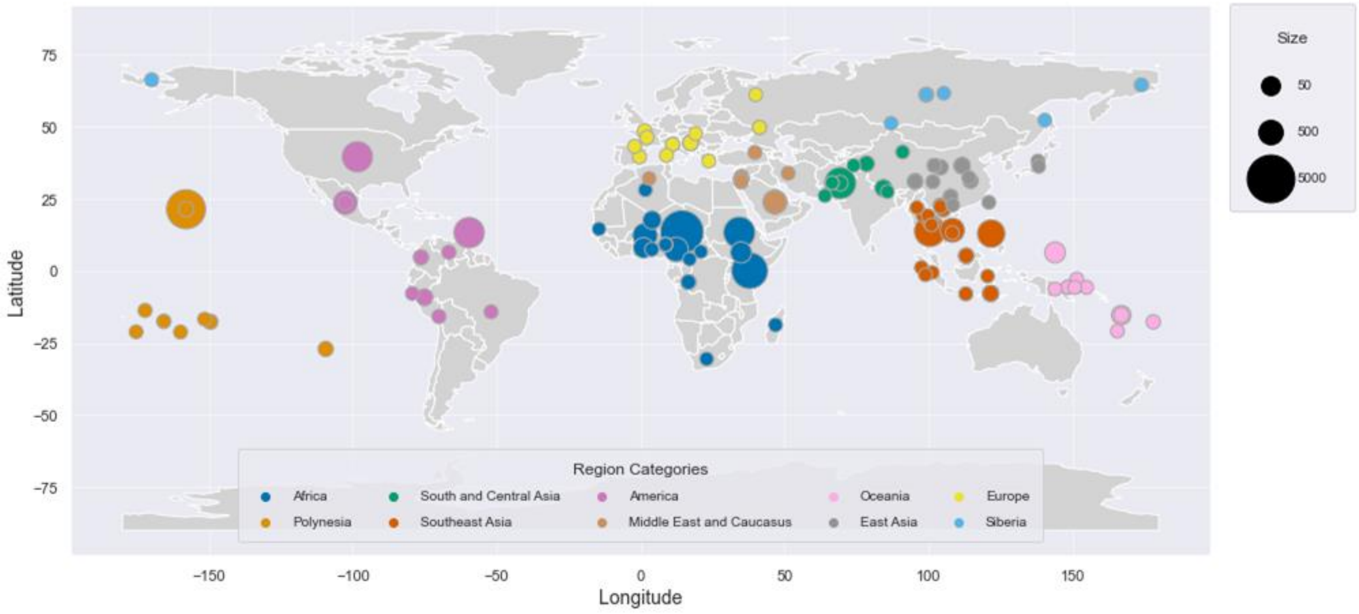


**Supplemental Figure 1: Regional distribution of Imputed Populations**. Each circle represents one population and the radius of the circle corresponds to the population size. The regional categories were assigned based on geographic location.

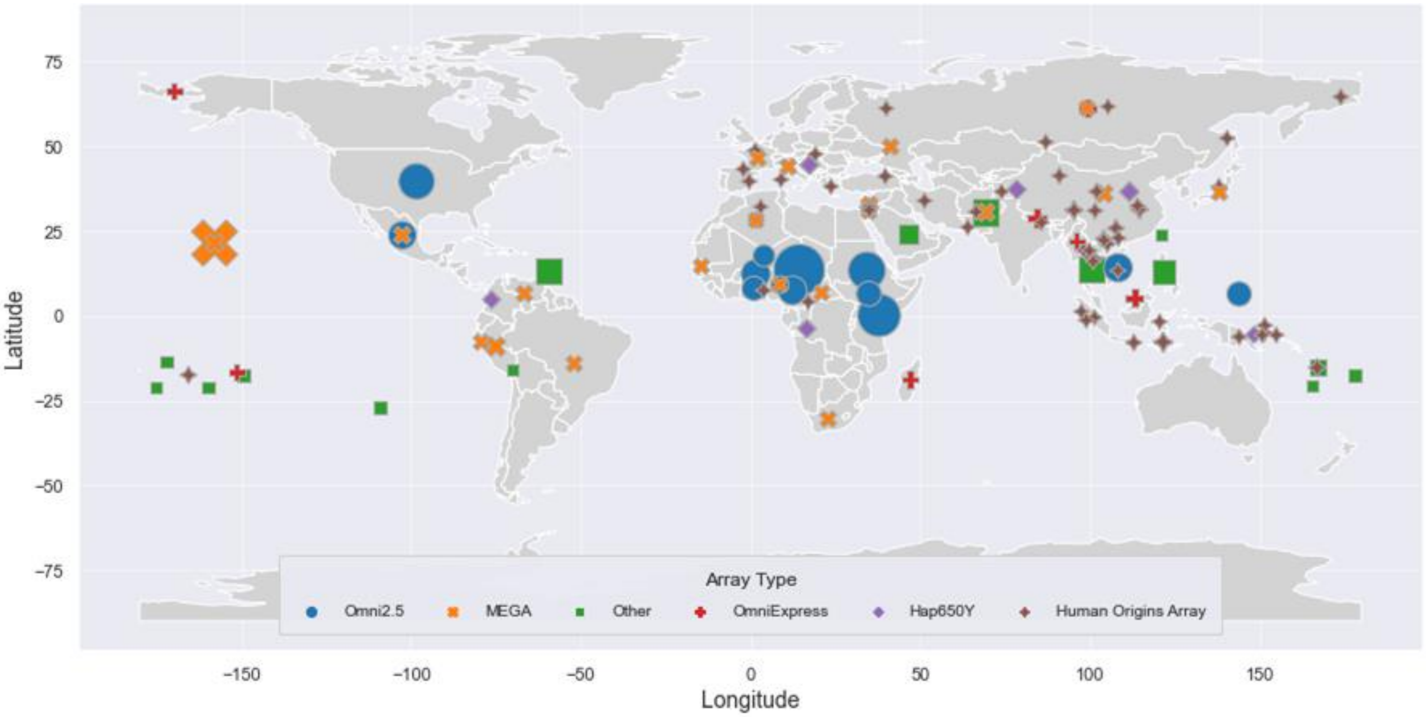


**Supplemental Figure 2: Distribution of Five Most Commonly Used Array Platform.** Each point represents one population and the radius of the circle corresponds to the population size. Only the top five most common arrays, Omni 2.5M, MEGA, and the Human Origins Array, HAP650Y, and the OmniExpress arrays are represented here. The remaining array types are designated as “Other”.

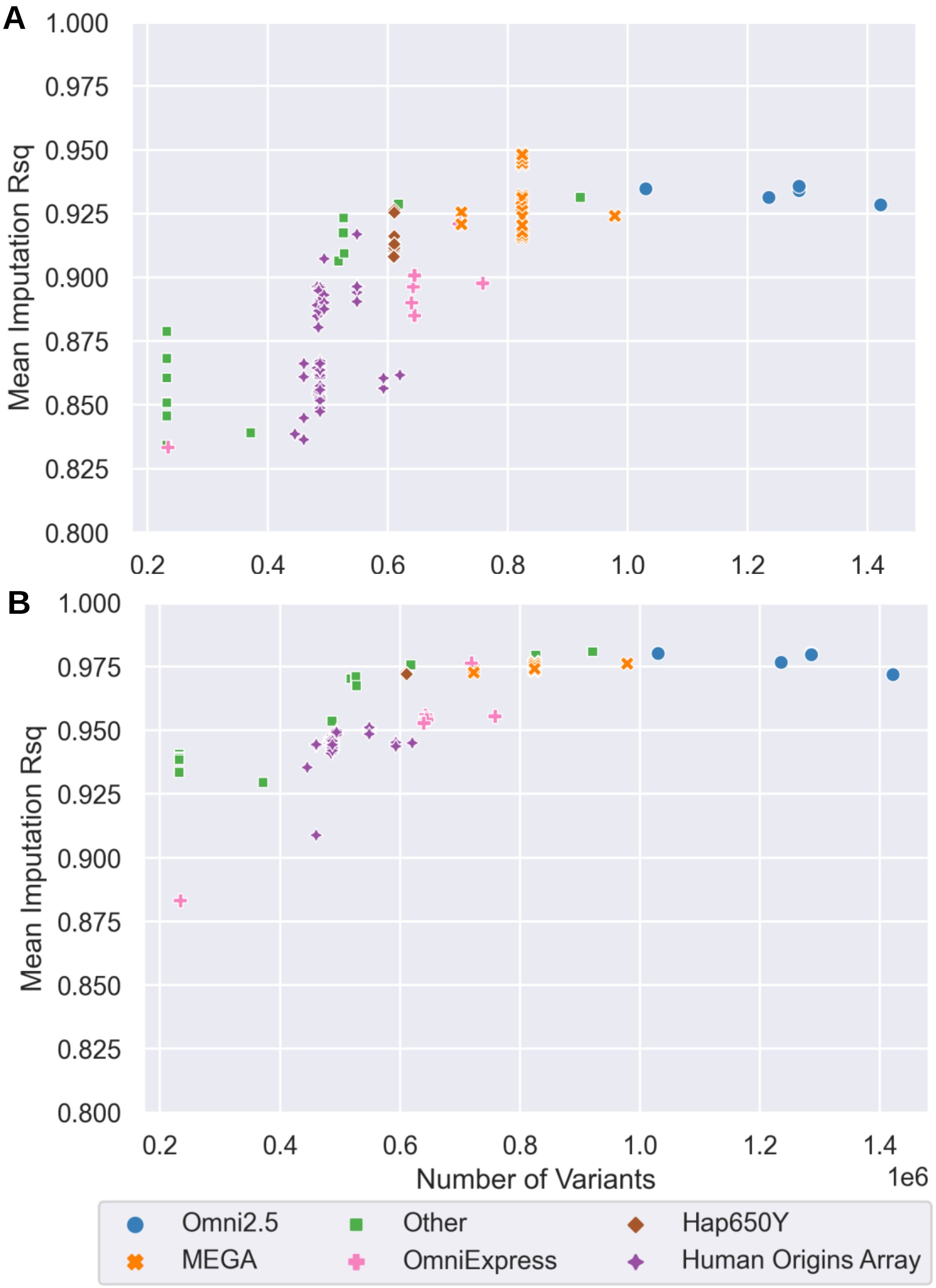


**Supplementary Figure 3: Imputation Quality as a Function of Marker Density.** Each point represents a subset of the UK10K cohort used in the primary analysis as the matched European control population. The points' shape and color denote the top five array types: Omni2.5, MEGA, Hapy650Y, OmniExpress, and the Human Origins Array. Remaining array platforms were collectively denoted as Other. Top panel (A) is for 1-5% alleles, bottom panel (B) is for 5-50% alleles. This figure shows a clear trend where high marker density is correlated with high average imputation Rsq.

**Supplemental Figure 4 (Separate PDF file): Imputation efficacy as measured by Rsq for each target population compared to matched European control population.** Stratified by minor allele frequency bins, each plot compared the distribution of imputation Rsq of the target population to that of a European cohort matched by sample size and SNP content.

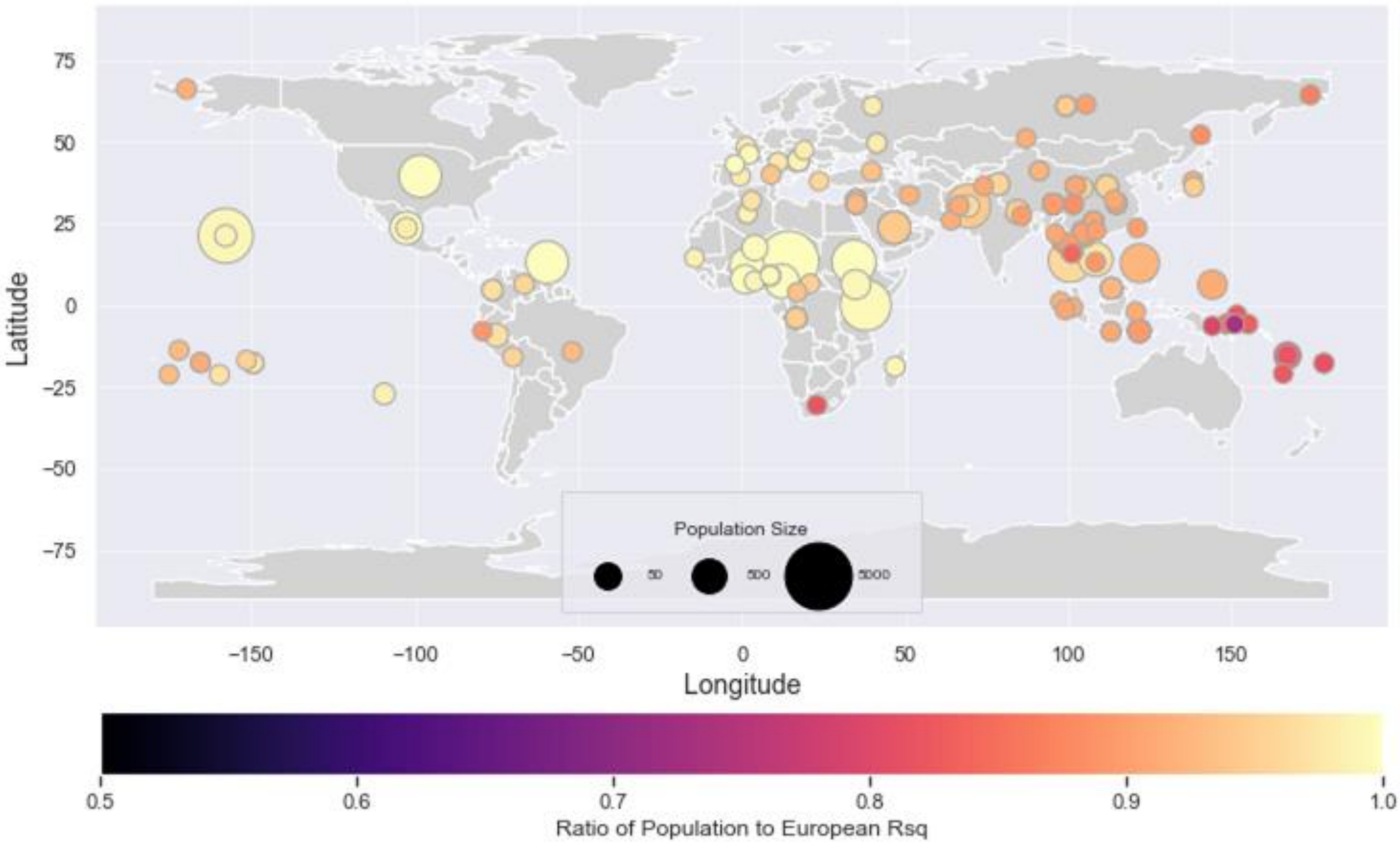


**Supplemental Figure 5: Ratio of mean Imputation Rsq of each population with UK10K controls for 5-50% alleles.** Values closer to 1.0 indicate no relative imputation quality loss compared to a European cohort matched by sample size and SNP content. Size of the circle scales with the sample size of the dataset that was imputed, and color of the circle scales with imputation quality.

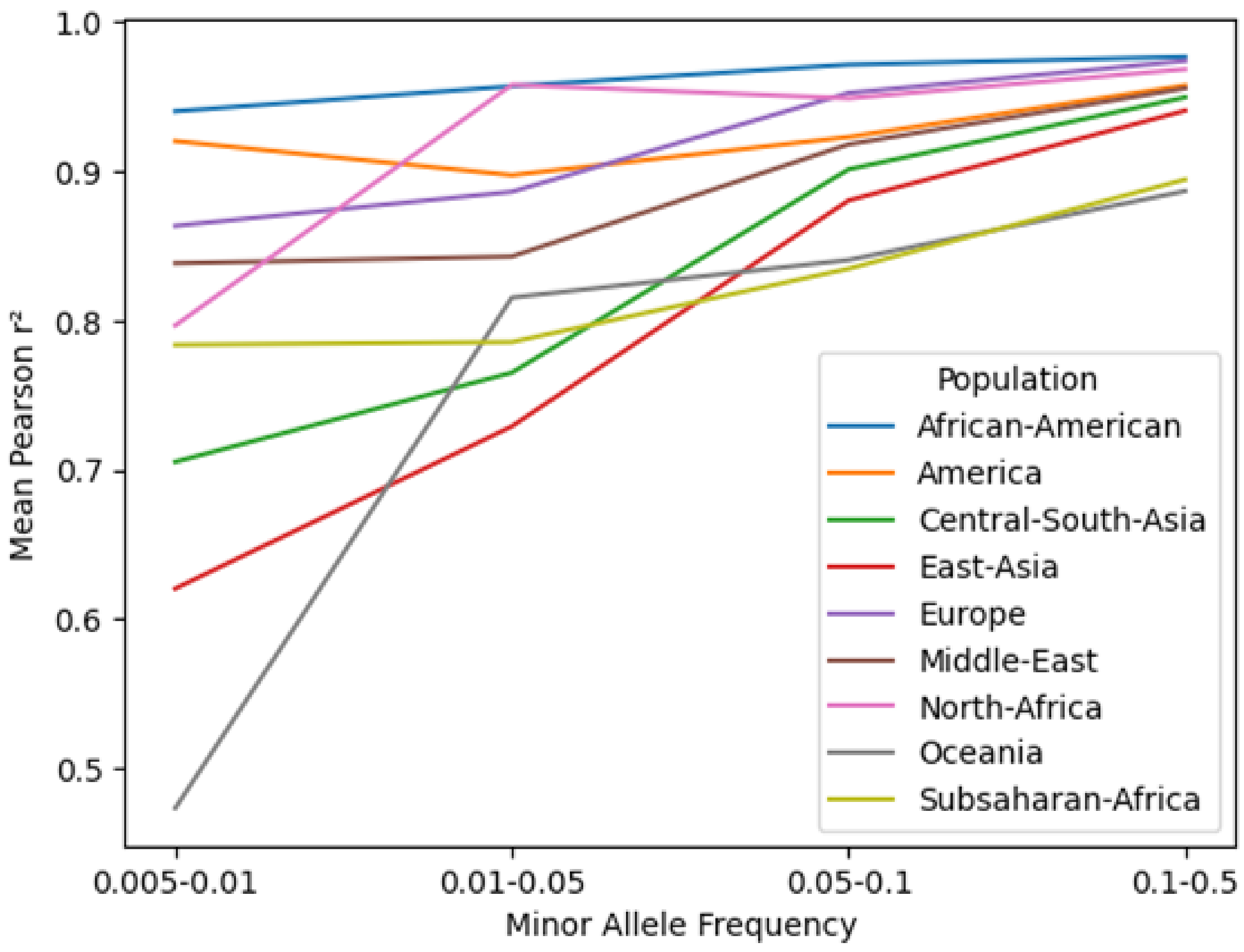


**Supplementary Figure 6: Average Pearson's $r^2$ for HGDP and AASP sequencing data**. All populations except for the African American population were sourced from the Human Genome Diversity Project. The African American population was from the African American Sequencing Project. These trends are generally consistent with the results using imputation Rsq. Note that the sample sizes are different across populations here (Supplemental Table 3), so the rarest allele frequency bin may not be interpretable for some populations with small sample sizes.

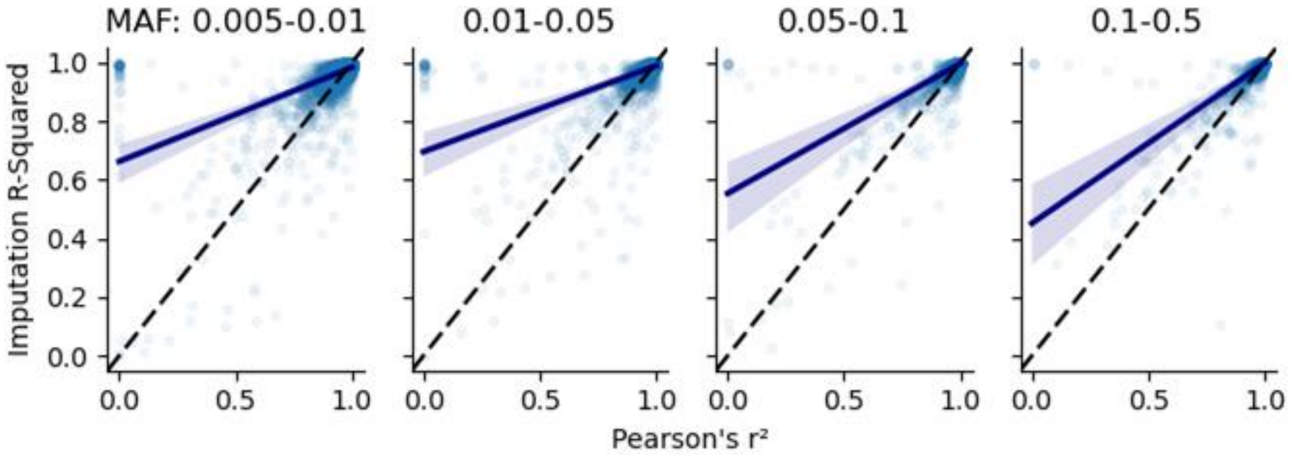


**Supplement Figure 7: The relationship between Pearson's r2 and Minimac4-Produced Imputation Rsq for individuals from the African American Sequencing Project.** A random set of 5,000 variants in each strata of minor allele frequency were sampled from 23andMe's African American Sequencing Project data (N=2269). The dash line denotes unity, while the solid lines are fitted linear models.

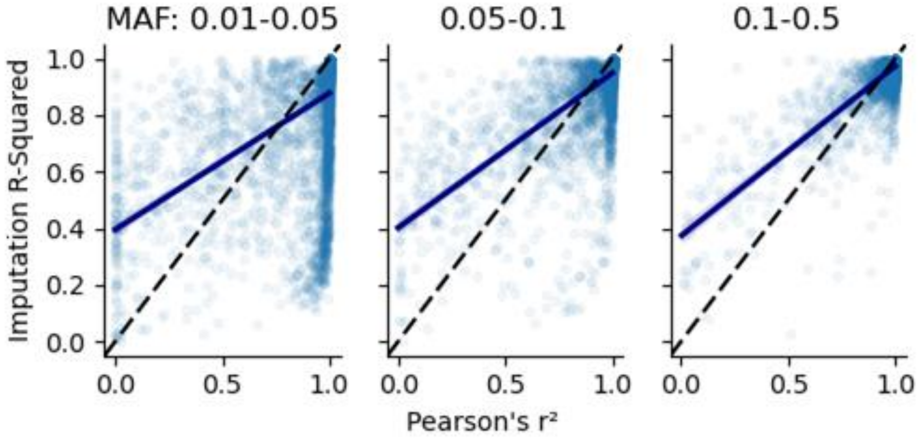


**Supplement Figure 8: The relationship between Pearson's r2 and Minimac4-Produced Imputation Rsq for individuals from the Americas (N=52) in the Human Genome Diversity Project.** A random set of 5,000 variants in each strata of minor allele frequency were sampled. The dash line denotes unity, while the solid lines are fitted linear models.

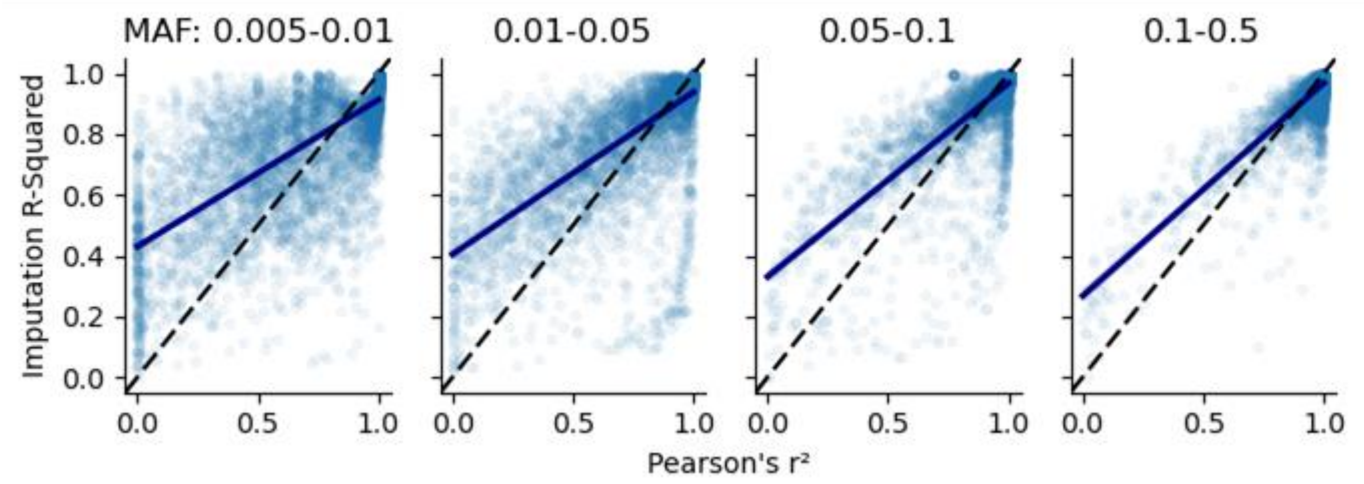


**Supplement Figure 9: The relationship between Pearson's r2 and Minimac4-Produced Imputation Rsq for individuals from South and Central Asia (N=195) in the Human Genome Diversity Project.** A random set of 5,000 variants in each strata of minor allele frequency were sampled. The dash line denotes unity, while the solid lines are fitted linear models.

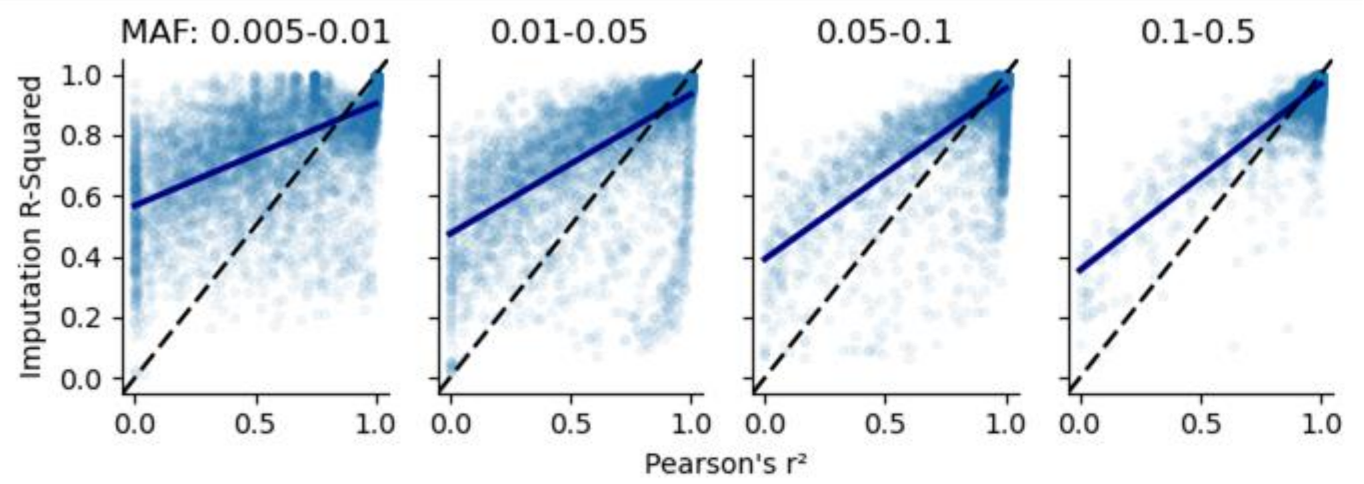


**Supplement Figure 10: The relationship between Pearson's r2 and Minimac4-Produced Imputation Rsq for individuals from East Asia (N=212) in the Human Genome Diversity Project.** A random set of 5,000 variants in each strata of minor allele frequency were sampled. The dash line denotes unity, while the solid lines are fitted linear models.

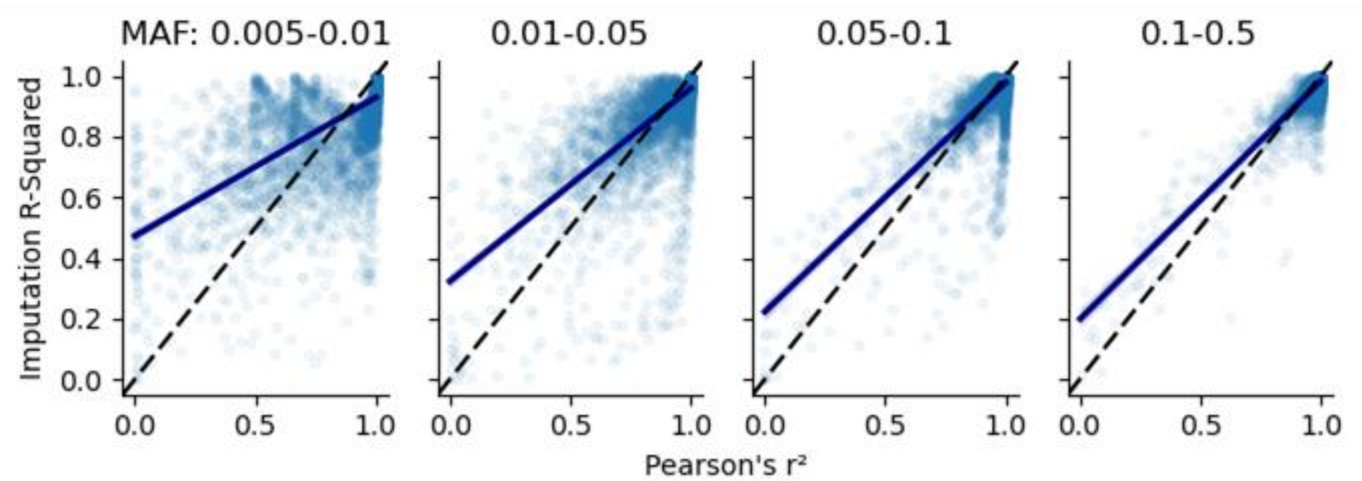


**Supplement Figure 11: The relationship between Pearson's r2 and Minimac4-Produced Imputation Rsq for individuals from Europe (N=145) in the Human Genome Diversity Project.** A random set of 5,000 variants in each strata of minor allele frequency were sampled. The dash line denotes unity, while the solid lines are fitted linear models.

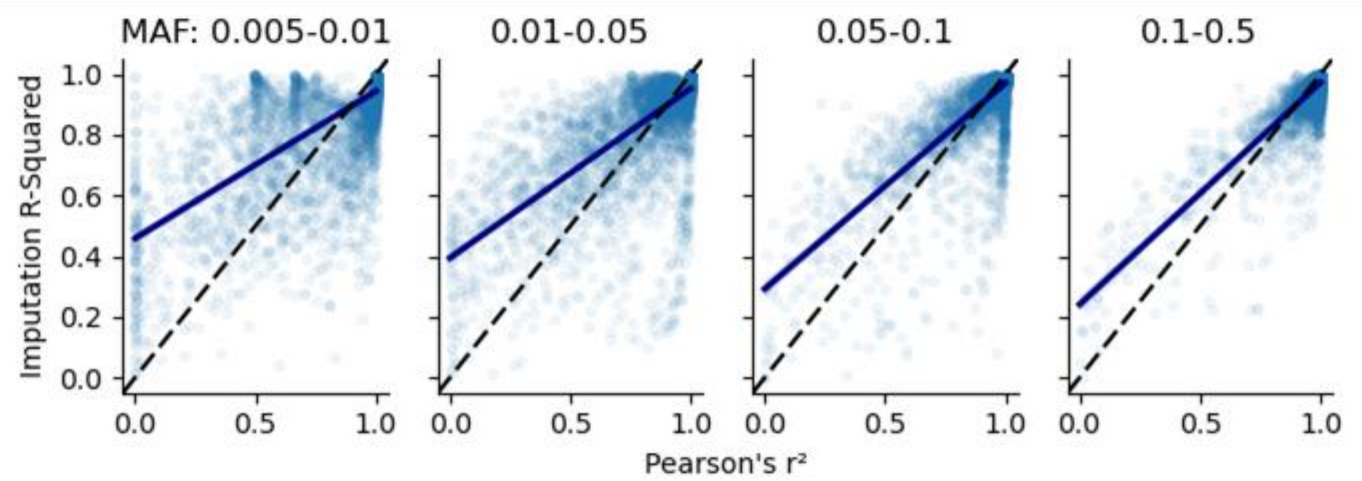


**Supplement Figure 12: The relationship between Pearson's r2 and Minimac4-Produced Imputation Rsq for individuals from Middle East (N=152) in the Human Genome Diversity Project.** A random set of 5,000 variants in each strata of minor allele frequency were sampled. The dash line denotes unity, while the solid lines are fitted linear models.

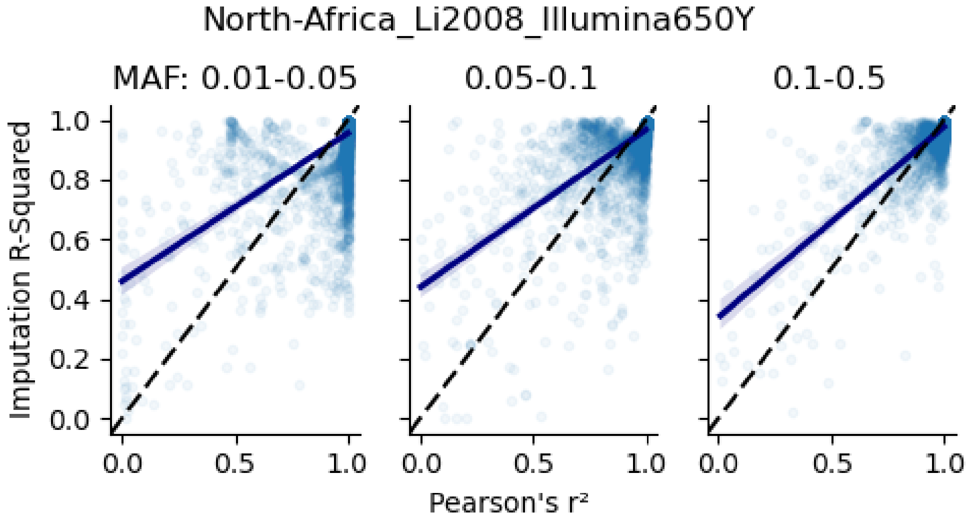


**Supplement Figure 13: The relationship between Pearson's r2 and Minimac4-Produced Imputation Rsq for individuals from North Africa (N=27) in the Human Genome Diversity Project.** A random set of 5,000 variants in each strata of minor allele frequency were sampled. The dash line denotes unity, while the solid lines are fitted linear models. The MAF bin for 0.005-0.01 were not analyzed due to small sample size in this population.

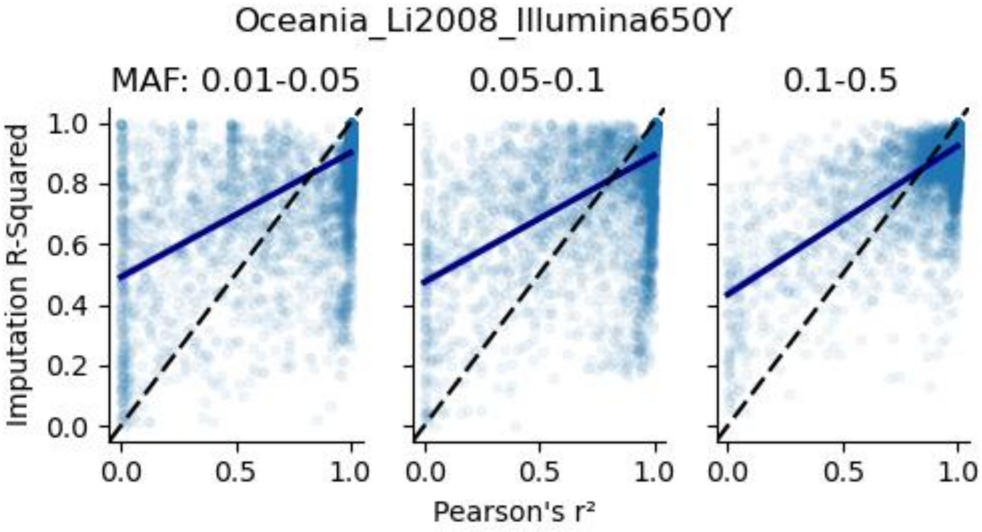


**Supplement Figure 14: The relationship between Pearson's r2 and Minimac4-Produced Imputation Rsq for individuals from Oceania (N=27) in the Human Genome Diversity Project.** A random set of 5,000 variants in each strata of minor allele frequency were sampled. The dash line denotes unity, while the solid lines are fitted linear models. The MAF bin for 0.005-0.01 were not analyzed due to small sample size in this population.

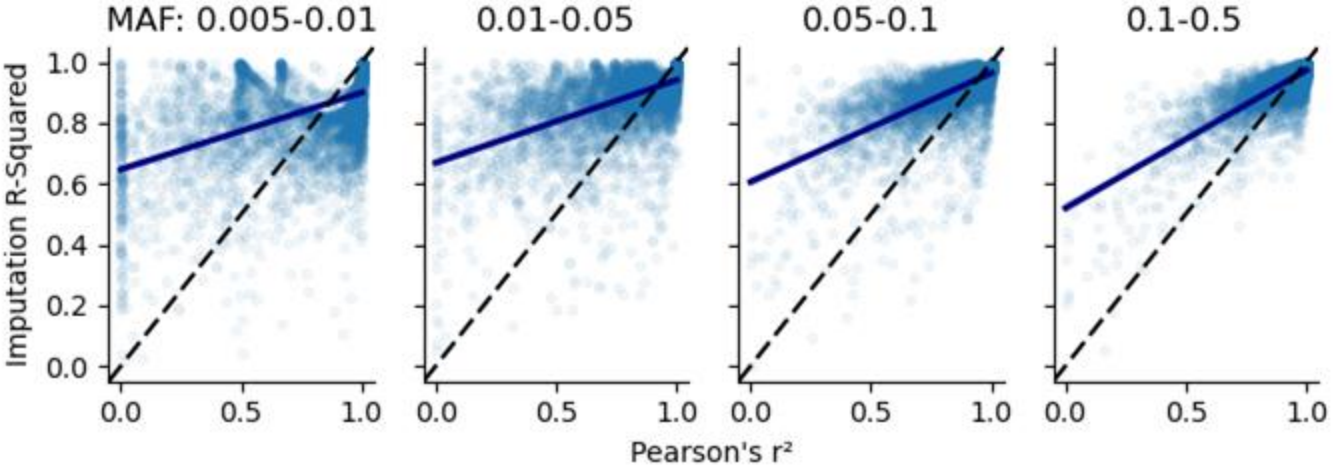


**Supplement Figure 15: The relationship between Pearson's r2 and Minimac4-Produced Imputation Rsq for individuals from Sub-Saharan Africa (N=100) in the Human Genome Diversity Project.** A random set of 5,000 variants in each strata of minor allele frequency were sampled. The dash line denotes unity, while the solid lines are fitted linear models.

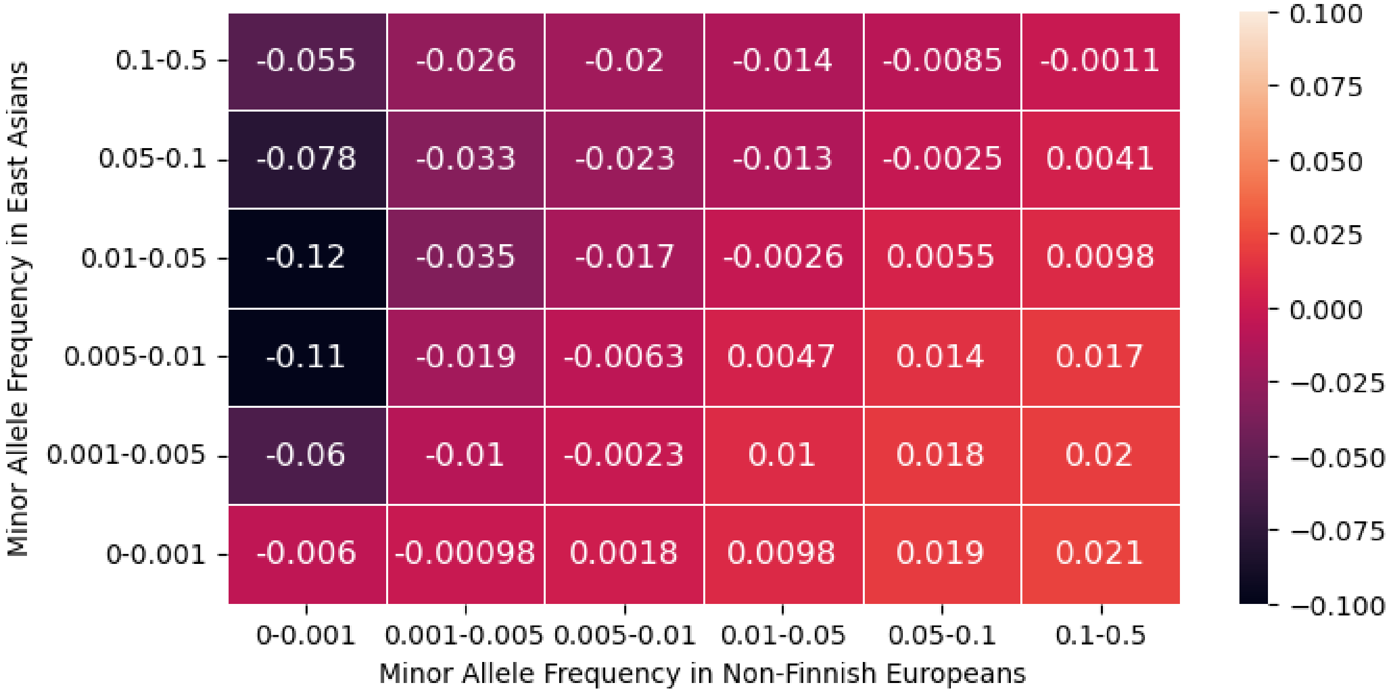


**Supplemental Figure 16: Difference in Imputation Quality between Meta-Imputation of TWB and TOPMed for Filipinos (N=1799) and TOPMed.** Stratified by minor allele frequencies estimated from non-Finnish European and East Asians from gnomAD, we computed the difference between imputation Rsq from meta-imputation (combining TOPMed and TWB imputation) with that from TOPMed imputation alone for a random sample of 10% of SNPs.

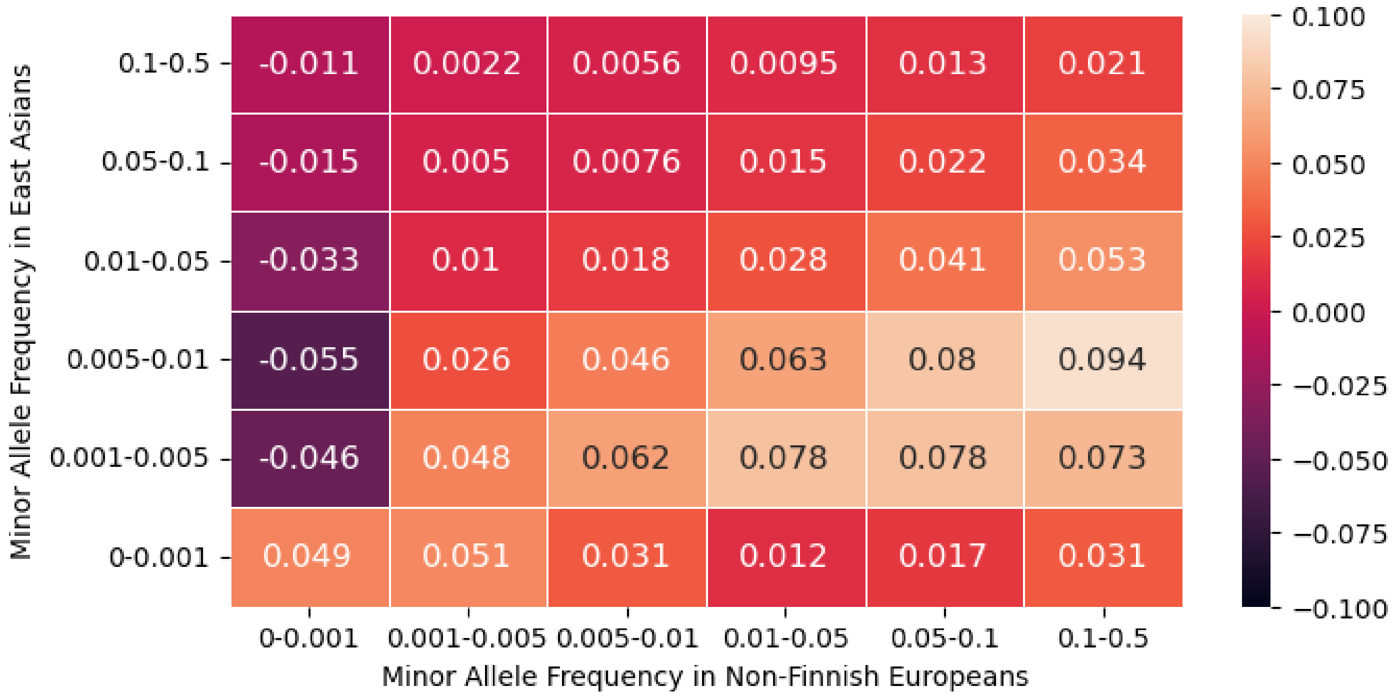


**Supplemental Figure 17: Difference in Imputation Quality between Meta-Imputation of TWB and TOPMed for Taiwanese (N=1999) and TOPMed.** Stratified by minor allele frequencies estimated from non-Finnish European and East Asians from gnomAD, we computed the difference between imputation Rsq from meta-imputation (combining TOPMed and TWB imputation) with that from TOPMed imputation alone for a random sample of 10% of SNPs.

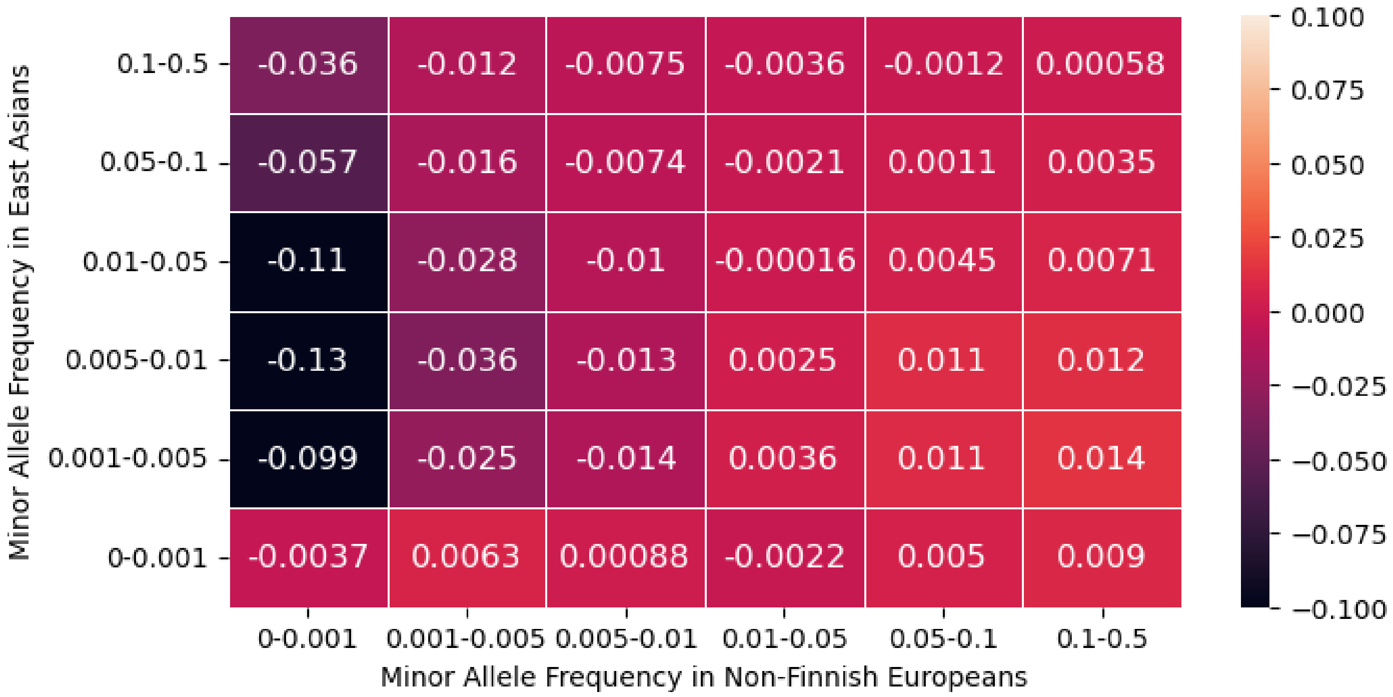


**Supplemental Figure 18: Difference in Imputation Quality between Meta-Imputation of TWB and TOPMed for Vietnamese (N=1264) and TOPMed.** Stratified by minor allele frequencies estimated from non-Finnish European and East Asians from gnomAD, we computed the difference between imputation Rsq from meta-imputation (combining TOPMed and TWB imputation) with that from TOPMed imputation alone for a random sample of 10% of SNPs.

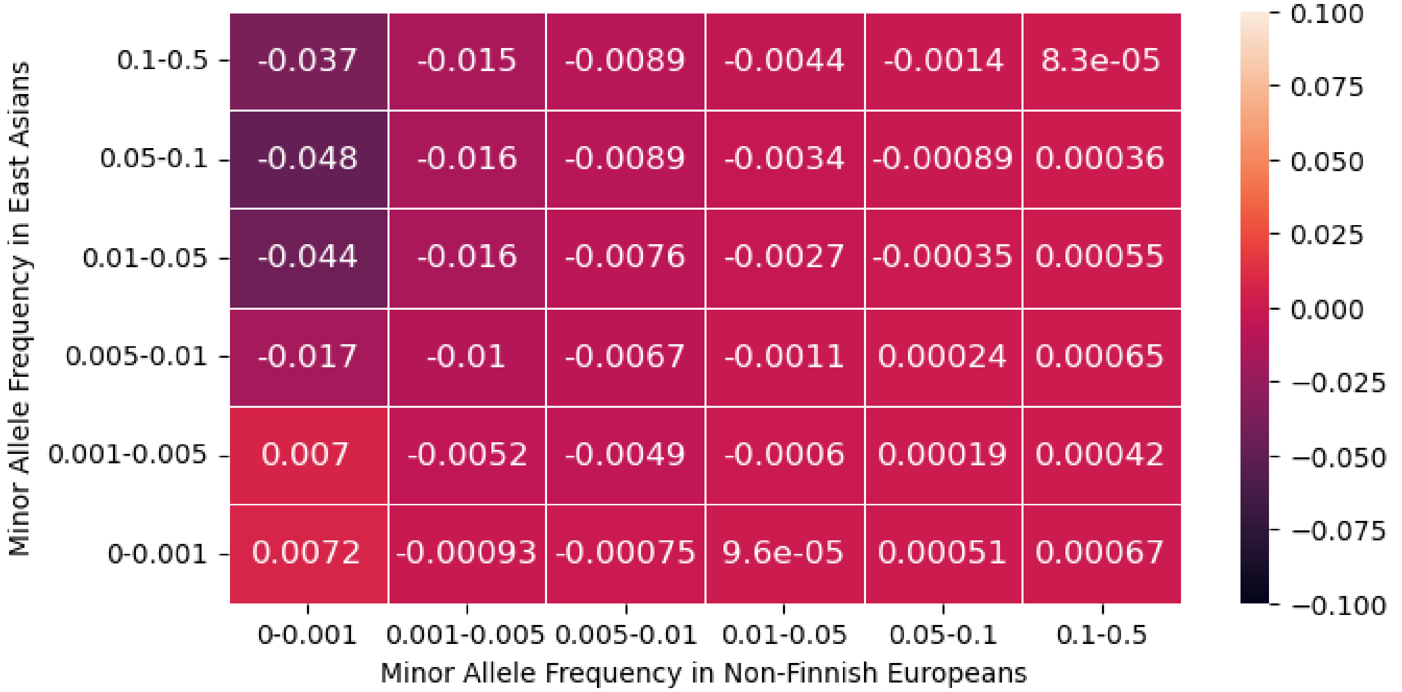


**Supplemental Figure 19: Difference in Imputation Quality between Meta-Imputation of TWB and TOPMed for Pakistani (N=2493) and TOPMed.** Stratified by minor allele frequencies estimated from non-Finnish European and East Asians from gnomAD, we computed the difference between imputation Rsq from meta-imputation (combining TOPMed and TWB imputation) with that from TOPMed imputation alone for a random sample of 10% of SNPs.

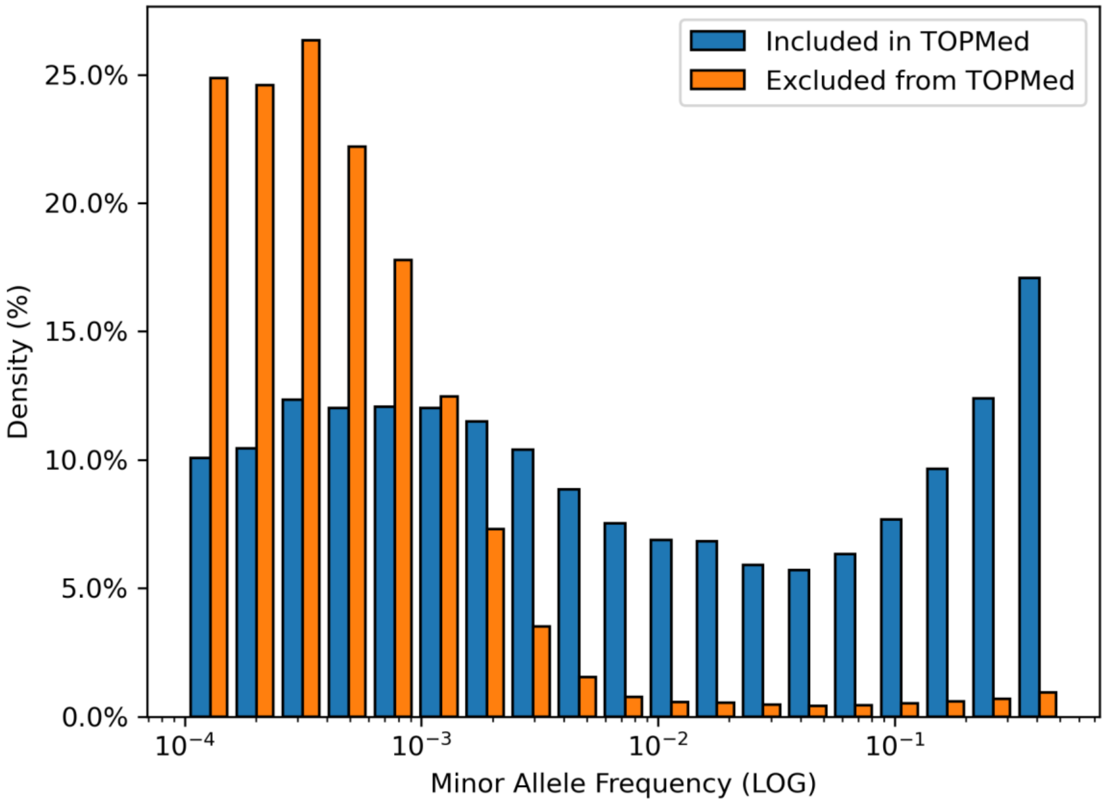


**Supplemental Figure 20: Variants found in TWB but excluded from TOPMed are rare in TWB.** The distribution of minor allele frequencies for a genome-wide random sample of 10% in TWB was plotted on a log scale and split between the variants included in both TOPMed (2,070,454) and TWB or in TWB only (1,923,111). Variants unique to TWB are predominantly rare (<0.001 MAF) in TWB.
